# Hijacking of the host cell Golgi by *Plasmodium* liver stage parasites

**DOI:** 10.1101/2020.07.24.220137

**Authors:** Mariana De Niz, Gesine Kaiser, Benoit Zuber, Won Do Heo, Volker T. Heussler, Carolina Agop-Nersesian

**Affiliations:** Institute of Cell Biology, University of Bern, CH-3012, Bern, Switzerland; Institute for Anatomy, University of Bern, CH-3012, Bern, Switzerland; Dept. of Biological Sciences, Korea Advanced Institute of Science and Technology (KAIST), Daejeon, Republic of Korea

**Author notes:** Present address: Instituto de Medicina Molecular João Lobo Antunes, Faculty of Medicine, University of Lisbon, Lisbon, Portugal (Mariana De Niz).

**Keywords:** host-parasite interaction, host cell Golgi, CLASP, Golgi-associated vesicular traffic, *Plasmodium* liver-stages, organelle hijacking

## Abstract

The intracellular lifestyle represents a challenge for the rapidly proliferating liver stage *Plasmodium* parasite. In order to scavenge host resources, *Plasmodium* has evolved the ability to target and manipulate host cell organelles. Using dynamic fluorescence-based imaging, we show a direct interplay between the pre-erythrocytic stages of *Plasmodium berghei* and the host cell Golgi during the entire liver stage development. Liver stage schizonts fragment the host cell Golgi into miniaturized stacks, which increases surface interactions with the parasite’s parasitophorous vacuole membrane. Interference with the host cell Golgi-linked vesicular machinery using specific dominant-negative Arf and Rab GTPases results in developmental arrest and diminished survival of liver stage parasites. Moreover, functional Rab11a is critical for the parasites ability to induce Golgi fragmentation. Altogether, we demonstrate that the structural and functional integrity of the host cell Golgi is necessary for optimal pre-erythrocytic development of *P. berghei*. The parasite hijacks the hepatocyte’s Golgi structure to optimize its own intracellular development.

## Introduction

Malaria is a mosquito-borne infectious disease caused by parasites of the genus *Plasmodium.* In the mammalian host, the liver stages of infection represent a bottleneck in the parasite’s life cycle. Hepatocyte infections are clinically silent, nevertheless, development of the parasite within the liver is extremely efficient, reportedly, one of the fastest growth rates among eukaryotic organisms (reviewed in (Prudencio et al., 2006)). To accomplish its extensive replication, the parasite depends not only on *de novo* synthesis of essential metabolites, but also on scavenging and storage of host-derived nutrients (Bano et al., 2007; Inácio et al., 2015; Itoe et al., 2014; Meis et al., 1984; Niklaus et al., 2019; Petersen et al., 2017; Sá E Cunha et al., 2017; Zuzarte-Luís et al., 2017).

A critical factor for a productive intracellular replication of *Plasmodium* is the ability to successfully form a parasitophorous vacuole (PV). The parasitophorous vacuole membrane (PVM) is the interface between the host cell cytosol, and the developing parasite. Although the role of the PVM and its molecular composition in liver stages is still not fully understood, this structure is thought to play a key role in nutrient acquisition (Bano et al., 2007; Fougère et al., 2016; Grützke et al., 2014; Petersen et al., 2017), waste disposal, and protection of the parasite from the host’s immune system (Agop-Nersesian et al., 2017; Boonhok et al., 2016; Grüring et al., 2012; Prado et al., 2015; Real et al., 2018; Spielmann et al., 2012). *Plasmodium,* like other intravacuolar pathogens, is believed to have evolved multiple strategies to access host cell resources. Among them, key elements include the generation of an export machinery, modification of PVM permeability to enable the uptake of cytosolic metabolites, and subversion of host cell organelles (reviewed in (Agop-Nersesian et al., 2018; Ingmundson et al., 2014; Spielmann et al., 2012)).

The Golgi apparatus is a key member of the secretory pathway, which structurally consists of multiple disc-like membranes that together form stacks with *cis*-to-*trans* polarity. Proteins received from the ER at the *cis*-Golgi are further processed and sorted for transport at the *trans-*Golgi network (TGN) from which they are destined to other intra- or extracellular locations. Aside of protein distribution, both the Golgi and the ER are major sites of lipid synthesis and glycosylation. Distinct processing, sorting, and synthesis events take place in an ordered sequence within different discrete regions of the Golgi complex (Reviewed in (Allan et al., 2002; Brandizzi and Barlowe, 2013; De Matteis and Luini, 2008; Lowe, 2011; Wilson et al., 2010). Compromise of the Golgi structure alters the overall function of this organelle. Several proteins are involved in the maintenance of the Golgi architecture, including members of the proteinaceous scaffold known as the Golgi matrix (GM), COPI, SNARE, and small GTPases including multiple members of the Rab protein family (Fig. 1 and Table 1). Various pathogens, including viruses, bacteria, and parasites, have been shown to interact with some of these proteins to scavenge nutrients and/or functionally or structurally alter the host cell Golgi (hereafter, hcGolgi) (Auer et al., 2019; Beske et al., 2007; Beyer et al., 2017; Cardoso et al., 2018; Cardoso et al., 2014; Coffey et al., 2015; Deffieu et al., 2019; Heuer et al., 2009; Lyu and Cai, 2019; Mañes et al., 2003; McGovern et al., 2018; Menger et al., 2002; Miller et al., 2017; Romano et al., 2017, 2013; Roy et al., 2006; Spanò and Galán, 2018; Spearman, 2018; Spriggs et al., 2019; Stein et al., 2012; Weber, Mary M. Faris, 2018).

**Figure 1.**
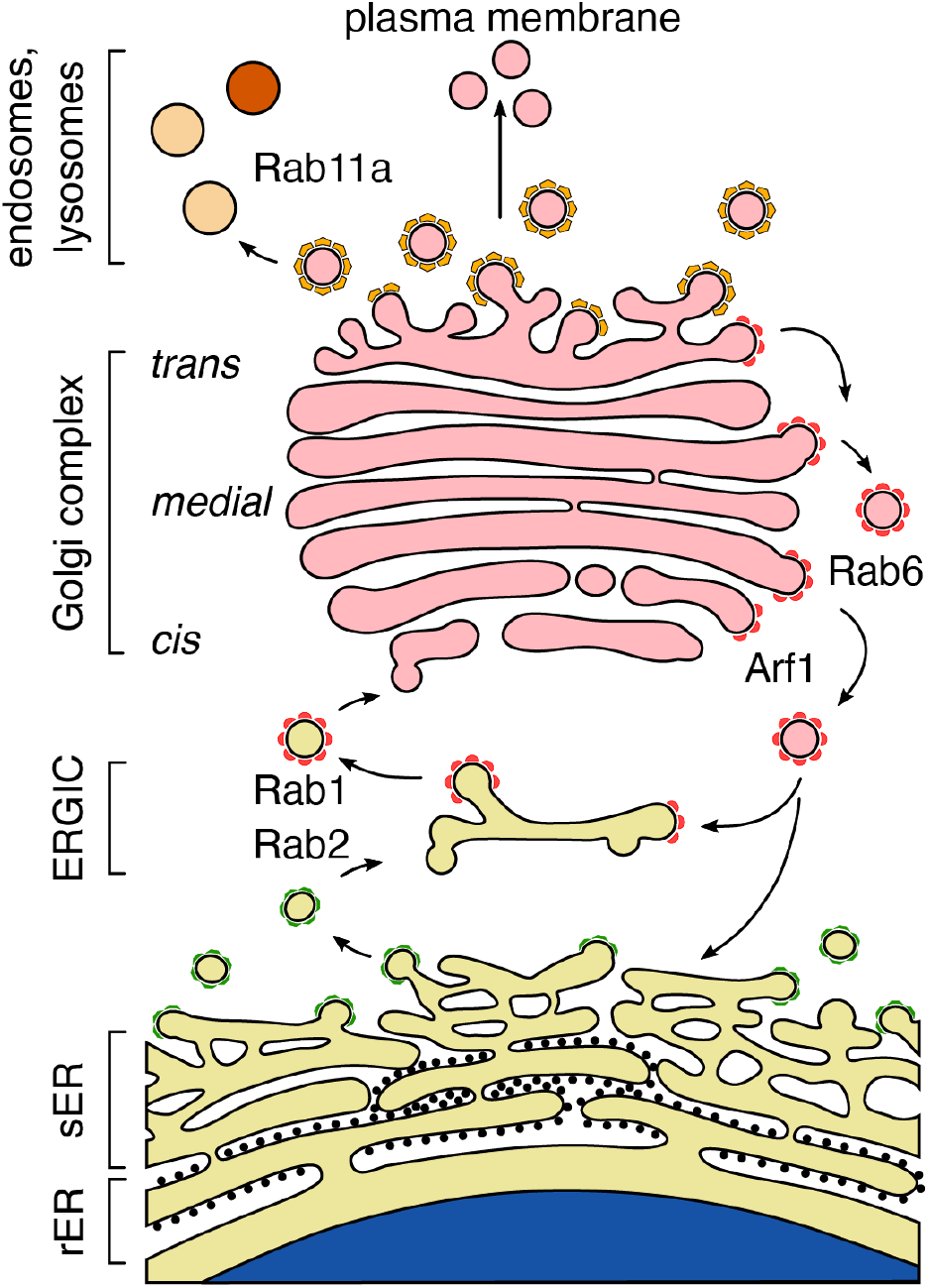
Simplified schematic of the endomembrane system in mammalian cells. Continued vesicle trafficking between the endoplasmic reticulum (ER) and the Golgi apparatus is important for Golgi biogenesis and maintenance. The two major sub-compartments of the ER are the rough ER (rER), composed of interconnected flattened sheets studded with ribosomes, and the entangled tubular network of the smooth ER (sER), where coatomer protein complex 2 (COPII) coated vesicles assemble at the ER exit sites (ERES) (anterograde transport shown on the left side of the schematic). COPII vesicles bridge the 300 to 500 nm space between the ER and the ER Golgi intermediate compartment (ERGIC), which is the first post-ER sorting compartment. Both Rab GTPases 1 and 2 coordinate the anterograde transport from the ER via the ERGIC towards the *cis*-Golgi. Rab1 regulates the post-ERGIC vesicle transport. There is evidence that COPI coated vesicles sustain the anterograde transport from the ERGIC to the *cis*-Golgi. The COPI vesicles play a well-established role in the retrograde intra-organelle traffic between the Golgi cisternae as well as for the transport from Golgi and ERGIC back to the ER (right side of the schematic). The COPI components are recruited by the ADP-ribosylated factor 1 (Arf1). At the *medial*-Golgi the retrograde travel of vesicle is mediated by the small GTPase Rab6. Based on the Golgi maturation model, the cisternae at the *cis* face of the Golgi mature to cisternae of the *trans*-Golgi network (TGN). At the *trans* face, the secretory side of the Golgi, Clathrin-coated vesicle bud off and are either exocytosed at the plasma membrane or targeted to the endo-lysosomal recycling compartment via Rab11.

**Table 1.**
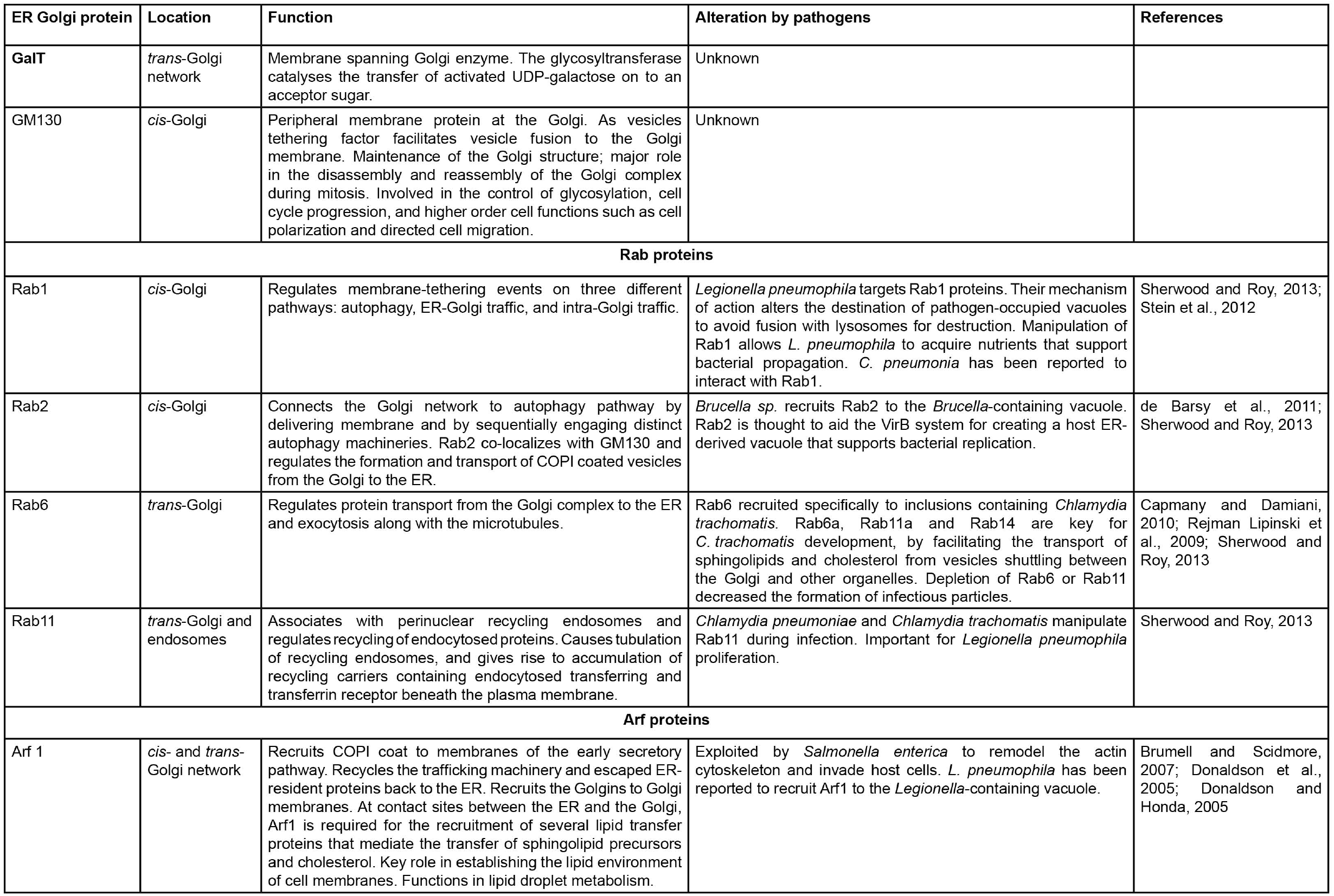
Golgi proteins studied in the context of Plasmodium exo-erythrocytic infections.

While PVM composition and nutrient incorporation is a topic of extensive research, *Plasmodium*-mediated subversion of host organelles is a less well studied area. Previously, Bano *et al.* showed that *Plasmodium* pre-erythrocytic stages settle at the juxtanuclear region of the host cell, and that throughout the development of the parasite, the PVM becomes associated with the host cell endoplasmic reticulum (ER) (Bano et al., 2007). While hcGolgi fragmentation was described, in-depth investigation of the relation of this organelle with *Plasmodium* was not undertaken. Our present study focuses on the interactions with, and recruitment of the hcGolgi by *P. berghei*.

In the present work, we explored the dynamics of interactions between the *P. berghei* PVM and the hcGolgi apparatus. We observe that pre-erythrocytic stages actively recruit the hcGolgi to the PVM, which acts as an interface between the parasite and the host. Mutant parasites with an impaired PVM function displayed deficiencies in hcGolgi recruitment, and disruption of hcGolgi structural integrity led to restricted parasite growth. Altogether, we determined that recruitment of and sustained contact with the hcGolgi, is key for the survival and development of the parasite.

## Results

### Recruitment of the host cell Golgi to the *Plasmodium berghei* PVM

Using quantitative fluorescence microscopy, we characterized the relation of *P. berghei* parasites to the hcGolgi throughout the entire pre-erythrocytic development. HeLa cells transiently transfected with the marker for the *trans*-Golgi network (TGN) the GFP-tagged β-1,4-galactosyltransferase (GalT-GFP), were infected with *P. berghei* ANKA sporozoites. The parasitophorous vacuolar membrane (PVM), in which the parasite develops, and its membranous expansion the tubovesicular network (TVN) was visualized by staining against either the early PVM proteins up-regulated in sporozoites 4 (UIS4), or exported protein 1 (EXP1). Laser scanning confocal (LSC) imaging of the proliferating liver-stage parasite revealed an intimate contact with the hcGolgi (Fig. 2A). To determine whether *Plasmodium*-hcGolgi contact occurs via the PVM or TVN, and whether this association changes as infection progresses, hcGolgi proximity to the PVM was quantified from 2 hours post-infection (hpi) onwards, up to 60 hpi (Fig. 2A-B). Already at 2 hpi, 45% of the invaded sporozoites were in contact with the hcGolgi (Fig. 2b). While a number of parasites in proximity to the hcGolgi showed association via the PVM (12%) (Fig. 2B left panel), the majority of contact events were via TVN protrusions (33%), bridging towards the extreme distantly located hcGolgi (Fig. 2B middle and lower panel). Moreover, hcGolgi proximity to, or contact with the parasite rapidly increased over the infection time. By 6 hpi, 78% of parasites were in contact with the hcGolgi. In the course of the early schizont’s development most parasites had established contact with the hcGolgi and maintained the relation with the PVM till the end of the liver stage development (Fig. 2B lower panel). Therefore, hcGolgi is recruited towards the parasite during the first 24 h of the infection. Additionally, we observe a shift from predominant TVN-association of the hcGolgi towards a PVM localization as infection progresses.

**Figure 2.**
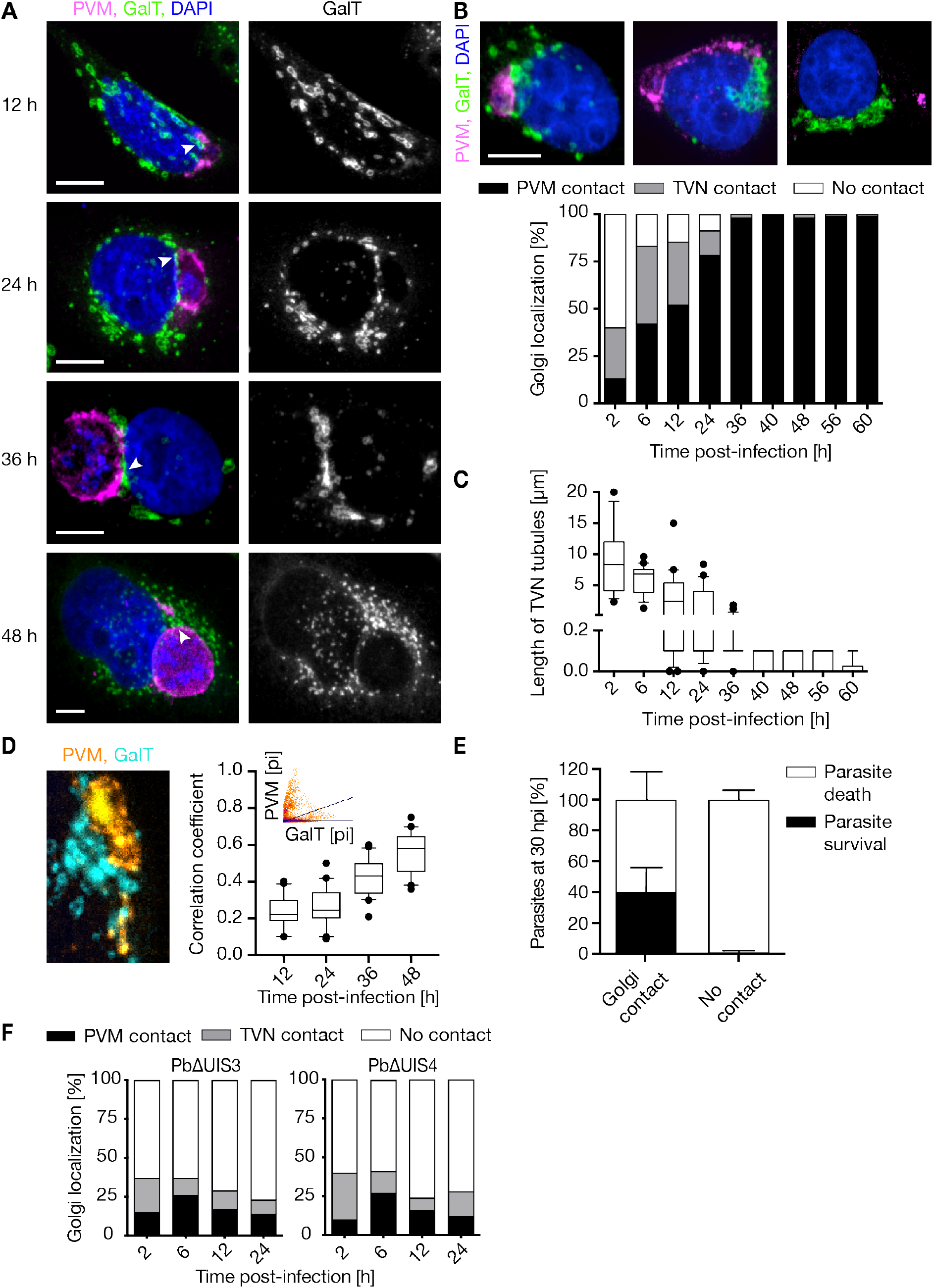
The parasite’s PVM and TVN establishes contact with the hcGolgi throughout infection. **A)** Confocal images of *P. berghei*-infected HeLa cells transiently expressing the *trans*-Golgi marker GalT-GFP (green) and co-stained with antibodies against PVM proteins UIS4 or EXP1 (magenta) and DAPI (blue). Representative images taken at progressing time points of the parasite development. Arrowheads indicate PVM/TVN contact with the hcGolgi. **B)** Quantification of the hcGolgi localization relative to the parasite across the entire liver-stage development. Top panel: Representative images of the hcGolgi-parasite interaction at 12 hpi. Images display GalT-GFP (green), PVM (magenta) and DAPI (blue). Classification of the interaction based on 1) contact the PVM (black), with GalT being in close proximity to the parasite body (left image); 2) contact via the TVN (grey), whereby the hcGolgi was located far away from the parasite body (beyond 5 μm), but contact was maintained by the parasite via the TVN tubular protrusion (middle image); 3) or no contact (white), whereby neither PVM nor TVN made contact with the hcGolgi (right image). **Lower panel:** Graph indicates the percentage of parasites displaying each type of localization relative to the hcGolgi at different time points of the infection. **C)** The length of the TVN extension reaching towards the hcGolgi was quantified at selected times post-infection. **D)** The increase in degree of co-localization between PVM (yellow) and GalT-GFP (turquoise) over the infection time. Co-localization was measured at the area of contact based on the Pearson’s correlation coefficient, by using the Coloc2 plugin available in Fiji, and confirmed with the co-localization threshold. **Left panel:** A representative image of co-localization is shown, together with the relevant scatterplot (inset on right panel). **Right panel:** Box plot displays the correlation coefficient of 20 images per time point. **E)** Quantification of the parasite death phenotype in liver schizonts (PbmCherry, 30 hpi) associated with or lacking hcGolgi contact. Parasite death was defined based on morphological features described by (Eickel et al., 2013). **F)** The importance of a functional PVM was evaluated by quantifying Golgi-association in PVM-deficient knockout parasites PbΔUIS3 (left graph) and PbΔUIS4 (right graph). Graph indicates the percentage of parasites displaying a hcGolgi-association (based on the classification described in **B)** during the first 24 h of infection. All data shown represent 3 independent experiments with n ≥ 100 infected cells per time point. Scale bars, 10 μm.

Quantification of the length of the TVN projections associated with the hcGolgi showed a progressive shortening of the TVN extension over time (Fig. 2C). While TVN extensions of sporozoites (2 hpi) display a median length of 8 μm and can reach up to 20 μm, from 12 hpi onwards this distribution becomes significantly reduced, resulting in an absolute minimum of 0.1 μm in mature schizonts (<40 hpi) (Fig. 2C). While hcGolgi contact via TVN extensions is rather characteristic for the early times post-infection, the PVM-hcGolgi contact becomes predominant at later stages.

Independent of the length of PVM or TVN extensions, the Pearson’s correlation coefficient for co-localization between the GalT-GFP and the parasite PVM, was measured around the contact site (Fig. 2D). Over time we observed a progressive increase in the degree of co-localization, indicating that more of the TGN becomes aligned on the surface of the PVM. Equally important, we observed that parasites without Golgi contact often display characteristic feature of parasite death (Eickel et al., 2013). We visually classified 30 h old PbmCherry liver schizonts depending on the association with the hcGolgi. Almost all parasites (99%) that lacked contact with hcGolgi presented morphological criteria of parasite death. When in contact with the hcGolgi, only 40-60% of the parasites displayed a death phenotype (Fig. 2E). Previously published live imaging results showed that in general only 50% of the liver stage parasites survive during the first 30 h of the develop (Prado et al., 2015).

Mutant parasites lacking the PVM protein UIS3 (PbΔUIS3) (Mueller et al., 2005b) or UIS4 (PbΔUIS4) (Mueller et al., 2005a) are surrounded by a PVM with impaired function. To visualize the PVM, mutants were stained with an anti-Exp1 antibody. Because both knockout (KO) lines fail to develop into mature liver stages, parasite numbers at all times after 24 hpi were minimal. Therefore, significant conclusions on the interaction are limited to 24 h. In the first 24 h of infection, WT parasites display the most pronounced increase in hcGolgi contacts with the PVM as described earlier (Fig. 2B). Although PbΔUIS3 and PbΔUIS4 showed a similar distribution regarding hcGolgi contact with PVM and TVN as WT parasites shortly after invasion (2 hpi), the number of contacts in the mutants did not increase over infection time. Both PbΔUIS3 and PbΔUIS4 showed significantly diminished hcGolgi contact via TVN protrusions and by adjacency with the PVM (Fig. 2F). Altogether, a compromised PVM function seems to have a direct impact on the PVM-hcGolgi interplay.

### Fragmentation of the hcGolgi leads to the formation of PVM-associated mini-stacks

Having determined the preferential localization of the parasite to the vicinity of the hcGolgi, we went on to determine changes in its morphology throughout *Plasmodium* infection. During LSC imaging of *P. berghei* at different time points of liver stage development, we noticed increased hcGolgi fragmentation, which was defined by a significantly wider spread of the hcGolgi, and a significantly higher number of Golgi stacks than in uninfected cells (Fig. 3A). In uninfected host cells the hcGolgi complex disassembles only when undergoing apoptosis or at the onset of mitosis to ensure correct partitioning into the daughter cells (Petrosyan, 2019). Quantification of hcGolgi fragmentation showed that at early times post-infection (6 to 12 hpi) up to 55% of infected cells exhibited a fragmented hcGolgi. At 24 hpi, this proportion increased to 85%, and continued to increase until 60 hpi, by which time hcGolgi fragmentation occurred in up to 99% of infected host cells (Fig. 3B). In uninfected host cells the proportion of fragmented hcGolgi remained constant at 12 ± 10% at all times. The proportion of hcGolgi fragmentation in infected cells correlated directly with the time and proximity of the hcGolgi to the parasite TVN and PVM (Fig. 3C). Golgi fragmentation is rather a consequence of parasite infection, than a simple result of mitotic disassembly.

**Figure 3.**
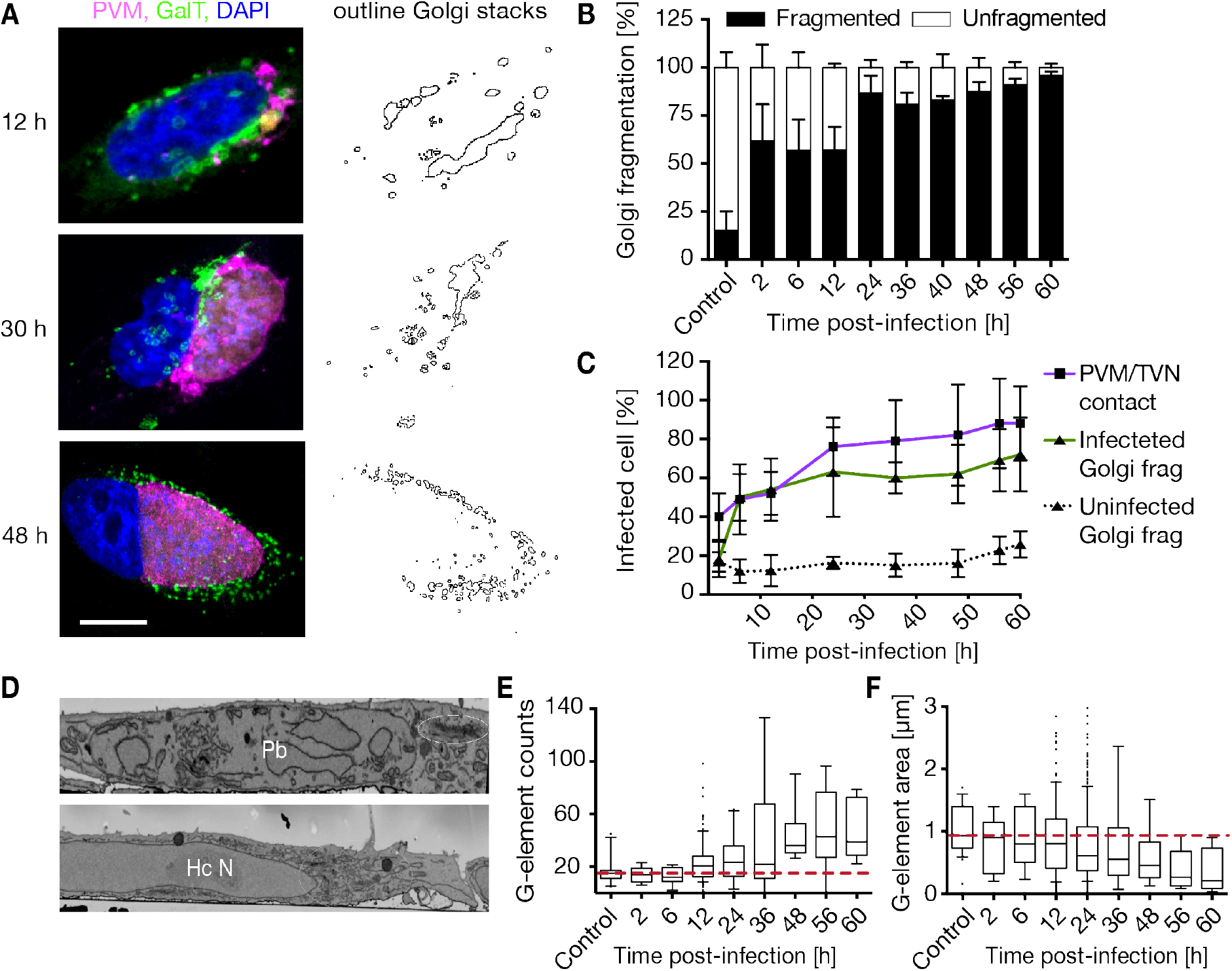
*P. berghei* liver infection promotes hcGolgi fragmentation. Infected cells transiently expressing GalT-GFP (green) were fixed at various times post-infection and stained for either UIS4 or EXP1 to visualize the PVM (magenta) and DAPI (blue). **A)** Immunofluorescence images of three representative time points during infection (12 hpi, 30 hpi and 48 hpi) highlight the progressing hcGolgi fragmentation. Right panel displays the bare outlines of the Golgi structure of images on the left. Golgi outlines were obtained using Fiji to schematically show stack distribution, number, and area covered. Scale bars, 10 μm. **B)** Quantification of the parasite-induced fragmentation of the hcGolgi. Percentage of infected cells that display hcGolgi fragmentation, as measured by stack distribution within the cells, and number of Golgi stacks. Uninfected cells were quantified to control for the effect of host cell transfection on Golgi structure and cell division mediated Golgi fragmentation. **C)** Relative percentages of host cells showing parasite PVM/TVN contact and hcGolgi fragmentation throughout infection. **D)** Serial block face scanning electron microscopy (SBF-SEM) on 48 h-infected cells (top panel) and uninfected control cells (lower panel) shows an morphologically intact Golgi apparatus. White circle indicate the hcGolgi. Labels: Pb, parasite; Hc N, host cell nucleus. **E-F)** Analysis of the degree of Golgi fragmentation during the course of infection. Quantification of Golgi elements (G-elements) numbers **(E)** and area **(F)** as infection progresses, showing an inverse trend. As infection progresses, the number of G-elements increase from > 20 to 70 per cell at 60 hpi. Equally, sizes of each G-elements decrease from an average of 1 μm^2^ to > 0.1 μm^2^. All data shown represent 3 independent experiments with n ≥ 200 infected cells per time point, quantified using Fiji.

To determine the ultrastructure of the hcGolgi fragments, we performed serial block face scanning electron microscopy (SBF-SEM) on 48 h-infected Hela cells. Uninfected host cells possess a very pronounced hcGolgi apparatus occupying a perinuclear position in close proximity to host cell ER. During *Plasmodium* infection, however, the hcGolgi is positioned adjacent to the PVM. Although the hcGolgi appears smaller in size, the parasite seems to preserve the typical hcGolgi architecture, consisting of a noticeable stack of cisternae (Fig. 3D). The parasite-dependent fragmentation of the hcGolgi seems to maintain the structural integrity and results in the formation of miniature organelles.

To analyse the degree of fragmentation over the infection period, we quantified total number and sizes (measured as areas μm^2^) of Golgi-elements (G-elements) in 3D-image reconstructions of uninfected and infected host cells (Fig. 3E-F). These G-elements are defined as GalT-GFP positive compartments and include different sized vesicles, individual or stacks of cisternae, or even the entire hcGolgi complex (a large G-element with a median of 1 μm^2^). During the first 12 h of infection, up to 50% of the infected host cells retained the perinuclear localization and ribbon-like morphology of the hcGolgi. This was the case despite the proximity of hcGolgi to the parasite. From 12 hpi, the number of G-elements compared to uninfected cells was almost 2-fold higher and continued to increase throughout infection. By 60 hpi, the number of G-elements was 4 to 7-fold higher than in uninfected controls (Fig. 3E). Conversely, while the area of each G-elements had a median of 0.9 μm^2^ in uninfected controls, sizes of elements gradually decreased throughout infection, becoming 10-fold smaller in host cells infected with parasites at 60 hpi (Fig. 3F). Altogether, host cells infected with *P. berghei* display a re-organisation and de-centralization of the hcGolgi.

### Host cell microtubule network supports hcGolgi-parasite interaction and parasite liver stage development

Microtubule (MT) networks play an important role in centralization of the Golgi at a perinuclear location, in maintaining a *cis-trans* polarity and Golgi ribbon assembly, as well as ensuring directional vesicle transport (de Forges et al., 2012; Wu and Akhmanova, 2017). To determine the contribution of host cell MT to the hcGolgi-PVM interactions, we decided to alter MT dynamics of infected cells. First, we explored the effect of two easily reversible MT inhibitors, namely Nocodazole, which binds to free tubulin heterodimers and accelerates depolymerization of MT, and Taxol, which binds to α-tubulin in the MT and stabilizes the MT polymers, protecting them from disassembly. The minimum concentration of Nocodazole or Taxol sufficient to elicit a disruptive effect on the hcGolgi apparatus was determined in uninfected cells by staining against alpha-tubulin and GM130 (Fig. S1A).

To determine whether host cell MT alter the localization and distribution of the hcGolgi around the parasite’s PVM, we treated HeLa cells infected with mCherry-expressing parasites with 50 nM of the respective drug for up to 3 h (Fig. 4A-C). We focused on 24 h parasites when host organelle-rearrangement becomes noticeably pronounced and parasite size did not significantly constrain the space of host cytoplasm. The degree of hcGolgi dispersion was evaluated based on GalT-GFP fluorescence in relation to the PVM and infected cells were subsequently classified into the categories 1) PVM-contact, 2) host cell cytoplasm and 3) host cell edge (Fig. 4A-B). In comparison to the untreated control parasites, drug-induced disassembly of the hcGolgi stack can be observed within the first 5 min after addition of either drug. After 30 min of drug incubation only 20% of the infected cells retained a prominent association of the hcGolgi with the PVM/TVN. Longer incubation times caused further scattering of the G-elements throughout the host cytoplasm resulting in a slight increase of infected cells with G-elements at the cell edge (Fig. 4A-B). When we quantified the fraction of G-elements that remained associated with the parasite in the inhibitor treated cell, we observed that 30% (Taxol) and 20% (Nocodazole) of the entire G-element population, did not dissociate from the PVM. Therefore, the parasite seems to form a stable, MT-independent connection with individual G-elements (Fig. 4C). Upon washout of the drugs, the G-elements reassembled within a maximum of 15 min into larger hcGolgi structures. In the majority of the infected host cells, this reassembly occurred directly at the vicinity of the parasite PVM (Fig. 4A-C).

**Figure 4.**
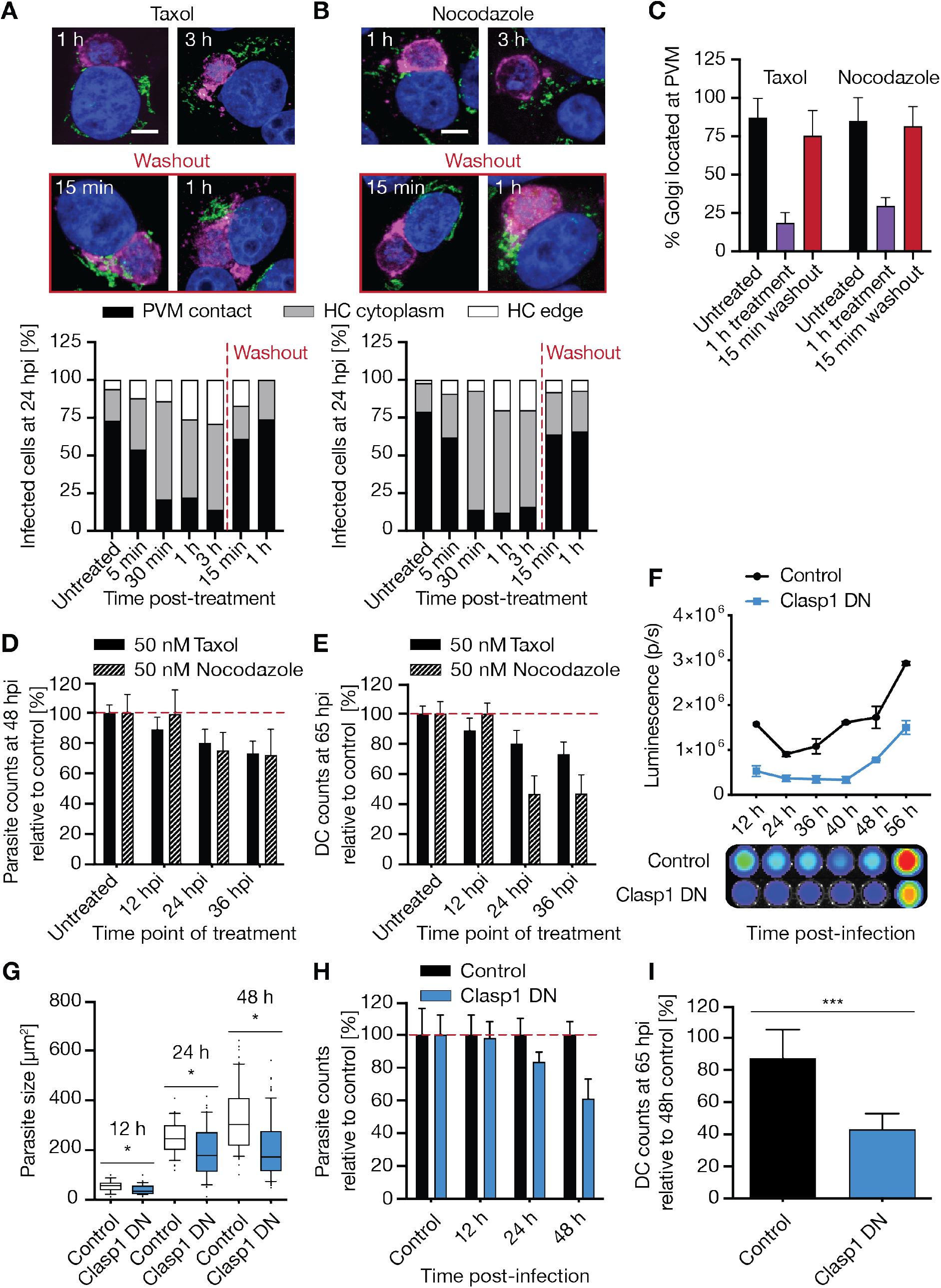
Interference with host cell microtubules (MT) affects parasite-hcGolgi interaction and pre-erythrocytic development of *P. berghei*. **A-B)** Effect of MT-acting drugs on the interconnection of the hcGolgi with the PVM. HeLa cells transiently transfected with *trans*-Golgi marker GalT-GFP (green) were infected with WT parasites. The 24 h old schizonts were treated with 50 nM of Taxol **(A)** or Nocodazole **(B)** as indicated. To allow MT lattice to recover, drugs were washed out from 3 h-treated infected cells, and hcGolgi positioning determined 15 min and 1 h after wash. The PVM in magenta was stained using anti-UIS4 or anti-EXP1 and nuclei in blue visualized with DNA dye, DAPI. Scale bar, 10 μm. **(A-B, lower panels)** Quantification of hcGolgi position following treatment with 50 nM Taxol **(A)** or 50 nM Nocodazole and after washout **(B)**. Depending on local maximum of the GalT-GFP signal in relation to the PVM, cells were classified according to 3 categories: 1) PVM contact, 2) host cytoplasm, and 3) cell edges. **C)** Percentage of hcGolgi that remains associated with the PVM in 24 h old schizonts following treatment with 50 nM Taxol or 50 nM Nocodazole and after recovery of the MT cytoskeleton. **D-E)** Impact of MT-dependent cell functions on the development of liver stage parasite. Infected cells were treated once for 1 h with 50 nM Taxol or 50 nM Nocodazole at different time points of parasite development. **D)** Quantification of total number of 48 h old schizonts relative to untreated cells, in percent. **E)** Percent reduction of detached cells shown relative to untreated cell at 65 hpi. **F-I)** Impact of non-centrosomal host cell microtubules on *P. berghei* liver-stage development. Transient overexpression of the microtubule binding domain of the CLIP-associating protein 1 (CLASP1), resulting in loss Golgi ribbon morphology and impaired intra- and post-Golgi cargo transport. **F)** Quantification of bioluminescence signals throughout the full pre-erythrocytic development of PbNLuc parasites in CLASP1 DN and GFP control cells. **(G-I)** Image-based quantification of the *P. berghei* liver stage development in CLASP1 DN or GFP expressing cells. **G)** Measurement of the parasite size during schizont proliferation. \**p* ≤ 0.05, paired *t*-test. **H)** Relative number of parasites compared to the GFP control shown in percent was determined at different times post-infection. **I)** Percent detached cells (65 hpi) shown relative to the number of parasites in GFP control cell determined at 48 hpi. All data shown corresponds to triplicate experiments and triplicate repeats per experiment, with n = 200 per cell type.

To evaluate the importance of the host cell MT cytoskeleton and its associated host cell functions (maintenance of hcGolgi architecture, spatial distribution of organelles and intracellular transport) on parasite proliferation, we went on to characterize parasite fitness after interfering with the MT network. Since treatment with Nocodazole and Taxol is reversible and enables us to distinguish which developmental stage is most vulnerable to MT manipulations, we treated infected cells once for 1 h at 12 hpi, 24 hpi or 36 hpi with either inhibitor. After careful washout of the drug, the parasites were further cultured to either allow development into 48 h schizonts (total parasite count) (Fig. 4D) or to complete liver stage development (detached cell formation) (Fig. 4E). While the 1 h inhibitor-treatment of young liver-stages (12 hpi) did not compromise parasite development, older 24 h and 36 h schizonts were sensitive in particularly to Nocodazole (Fig. 4D-E). However, with this experimental setup we could not exclude that both drugs might also negatively affect the parasite’s MT.

To eliminate this potential confounder, we transiently overexpressed the microtubule binding domain of the CLIP-associating protein 1 (CLASP1) in HeLa cells, resulting in a dominant negative phenotype (Maiato et al., 2003a, 2003b). CLASP1 is a microtubule plus-end tracking protein that promotes the stabilization of noncentrosome-derived microtubules. It is essential for the control of Golgi-derived microtubules keeping ribbon morphology, and the intra- and post-Golgi cargo transport alive (Miller et al., 2009). Transient transfection of host cells with EGFP-CLASP1 DN resulted in an extensive dispersal of the hcGolgi in transfected cells (Fig S1B). Subsequent infection with a transgenic PbNLuc *P. berghei* line constitutively expressing NanoLuc luciferase (De Niz et al., 2016) showed a significant decrease of luminescence during the proliferative phase of the parasite (12 hpi onwards) (Fig. 4F). Our imaging-based quantitative analysis revealed that the phenotype was due to a reduction in parasite sizes as well as overall parasite numbers (Fig. 4G-H). Furthermore, the loss of CLASP1-mediated Golgi organization and function prevents efficient parasite development and survival, as assessed by detached cell formation (Fig. 4I).

### Identification of Golgi-associated small GTPases with negative impact on the parasite fitness

Since a direct interaction of the hcGolgi with the PVM appears to support progression and survival of the exo-erythrocytic forms (EEF), we further dissected the importance of the cellular function of the hcGolgi. To subvert hcGolgi maintenance and organization, we interfered with Golgi-associated vesicular traffic by expressing distinct sets of dominant negative GTPases. The effect on parasite development was quantified by bioluminescence and fluorescence imaging. We selected five candidates to explore the effects of overexpression of dominant negative mutants on *Plasmodium* development. These candidates were the ADP-ribosylation factor (Arf1) – a target of Brefeldin A – that recruits COPI complex and regulates ER-Golgi and intra-Golgi vesicle transport (Beck et al., 2009; D’Souza-Schorey and Chavrier, 2006; Gillingham and Munro, 2007; Kawasaki et al., 2005), and four members of the small Rab GTPase family Rab1a, Rab2, Rab6a, and Rab11a that orchestrate the vesicular traffic at three main locations of Golgi complex. The small GTP-binding proteins Arf1, Rab1a- and Rab2-regulate the vesicle transport between ER-exit sites and the *cis*-Golgi. Dominant negative (DN) mutants of the two ER-Golgi Rabs (1a and 2) as well as Arf1 mutant, with the single threonine exchanged to asparagine at position 31 (T31N), cause disassembly of the hcGolgi. The effect on hcGolgi architecture has also major implications on its cellular function. At the *medial*-Golgi, Rab6 mediates retrograde intra-organelle traffic between the Golgi cisternae, while Rab11a is associated with the TGN. Importantly, loss of function mutations for Rab6 and Rab11a only mildly interfere with the structural organisation of the Golgi (Fig. 1 and Table 1) (Goud et al., 2018; Kjos et al., 2018; Pfeffer, 2017; Short et al., 2005). As an initial assessment, we monitored parasite (PbNluc) load in HeLa cells ectopically expressing the dominant negative (DN) mutants of Arf1, Rab1a, Rab2, Rab6a or Rab11a, by bioluminescence (Fig. 5 and Fig. S2). Normal parasite growth was assessed in HeLa cells transiently expressing a cytoplasmic GFP reporter (control). The inactive Arf1 mutant had the most severe effect on the entire parasite development. Already at 12 hpi parasite luminescence was reduced by 50% and did not recover during the ongoing infection. When blocking anterograde ER to Golgi vesicle traffic with Rab1aDN (Reviewed in Goud et al., 2018), a 40% decrease in luminescence signal was observed at 36 hpi, which continues to further decline. A similar trend was observed with Rab11a (40% reduction at 40 hpi), which regulates the exocytic (trans-Golgi to the plasma membrane) and endocytic recycling pathway (Reviewed in (Goud et al., 2018; Welz et al., 2014)). Both Rab2DN and Rab6aDN resulted only a slight developmental delay during the first 24 h. However, at later time points parasites were able to recover, showing no significant impact on the overall parasite fitness (Fig. 5B). Therefore, we did not proceed with further characterisations of the Rab2a and Rab6a mutants.

**Figure 5.**
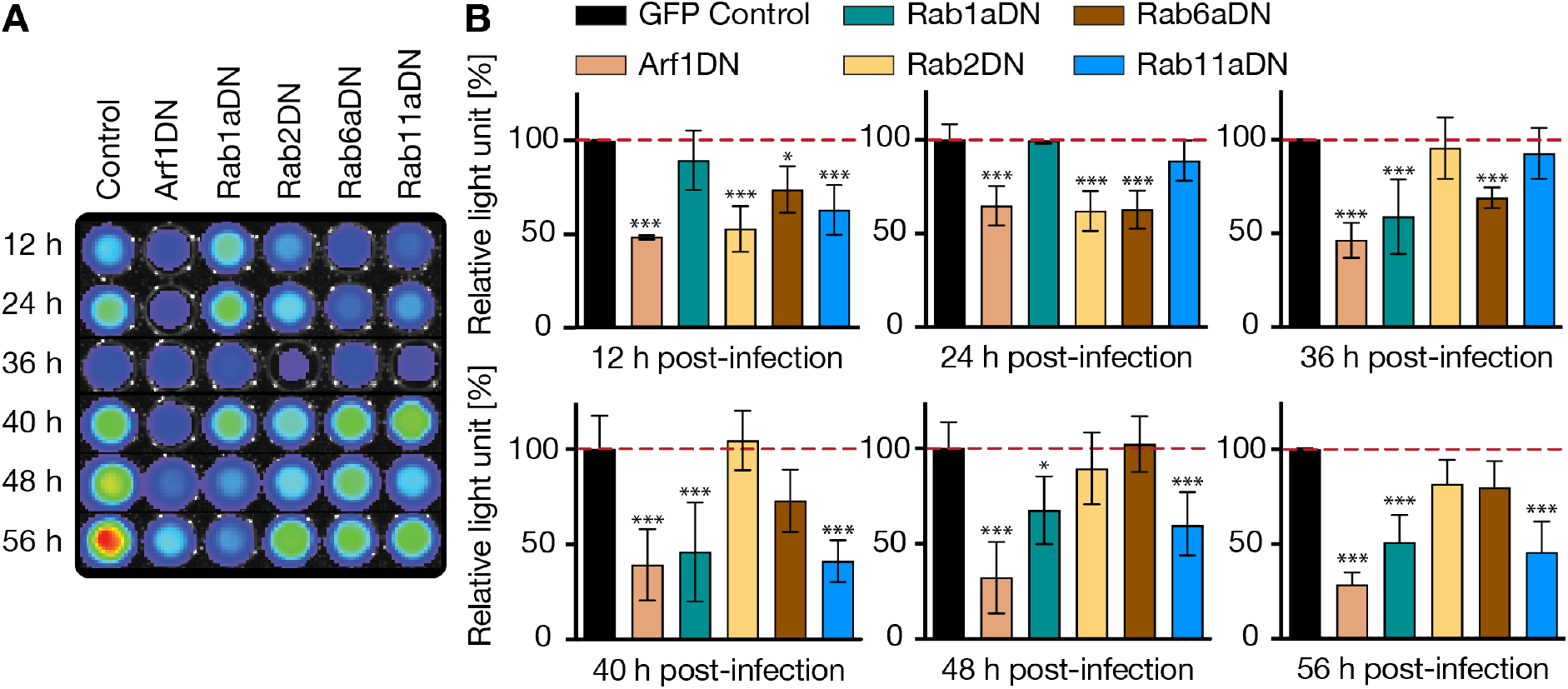
Bioluminescence based targeted screening for regulators of Golgi-associated vesicular transport that hamper the fitness of *P. berghei* liver stage. HeLa cells transiently overexpressing either GFP (control), or one of the dominant negative GTPase (Arf1DN, Rab1aDN, Rab2DN, Rab6aDN and Rab11aDN) were infected with NanoLuc-expressing *P. berghei* parasites (PbNLuc). **A)** Cells were lysed at various times post-infection (12 h, 24 h, 36 h, 40 h, 48 h and 56 h) and bioluminescence of each set of cells measured using an IVIS Lumina II system. **B)** Quantitative representation of luminescence measurements at every given time during infection. Parasite luminescence at each time point is represented relative to luminescence in GFP control, represented as 100%. All data are shown as the mean ± SD from 3 independent experiments with duplicate samples per experiment. *p ≤ 0.05 and ***p ≤ 0,01, paired t-test.

### Cellular function and structural integrity of the hcGolgi is important for*P. berghei* development

Based on our initial luminescence-based screening, we decided to further dissect the developmental phenotype of the parasites and also analyse the parasite-induced hcGolgi-fragmentation in cells deficient for Arf1-, Rab1a- and Rab11a-dependent vesicle transport. For the imaged-based quantifications, PbmCherry-infected HeLa cells were fixed at various time points of development. The changes in Golgi morphology and subcellular location induced by the loss of GTPase activity (Arf1, Rab1a or Rab11a) and by the parasite infection were visualized by staining against the *cis*-matrix protein GM130.

The two small GTPases, Arf1 and Rab1a, tightly regulate opposite vesicle routes between the ER and Golgi (Fig. 1). In both dominant negative mutants the loss of function results in a fast and complete disassembly of the hcGolgi complex and a loss of hcGolgi homeostasis (Fig. S3), confirming observations previously shown in other contexts (Donaldson et al., 2005; Dascher et al., 1994). After infection, we compared the number of G-elements between cells expressing either GFP or Rab1aWT as controls, and the respective DN mutant (Arf1DN or Rab1aDN) cells. In host cells infected with 12 h to 48 h old schizonts, the G-element counts were significantly increased in both mutants. While infected control cells display the characteristic parasite-mediated hcGolgi fragmentation, we detect 2x more G-elements in the Arf1DN and 1.5x more in Rab1aDN deficient cells already at 12 hpi. The phenotype becomes even more pronounced in late schizonts (Fig. 6A, left and middle panel). This enhanced disintegration of the hcGolgi suggests a cumulative effect triggered by the parasite-induced hcGolgi fragmentation and DN-dependent structural loss of the hcGolgi. Because both mutants display the characteristic hcGolgi disassembly also in presence of *P. berghei* infection, we anticipate a loss of its cellular function.

**Figure 6.**
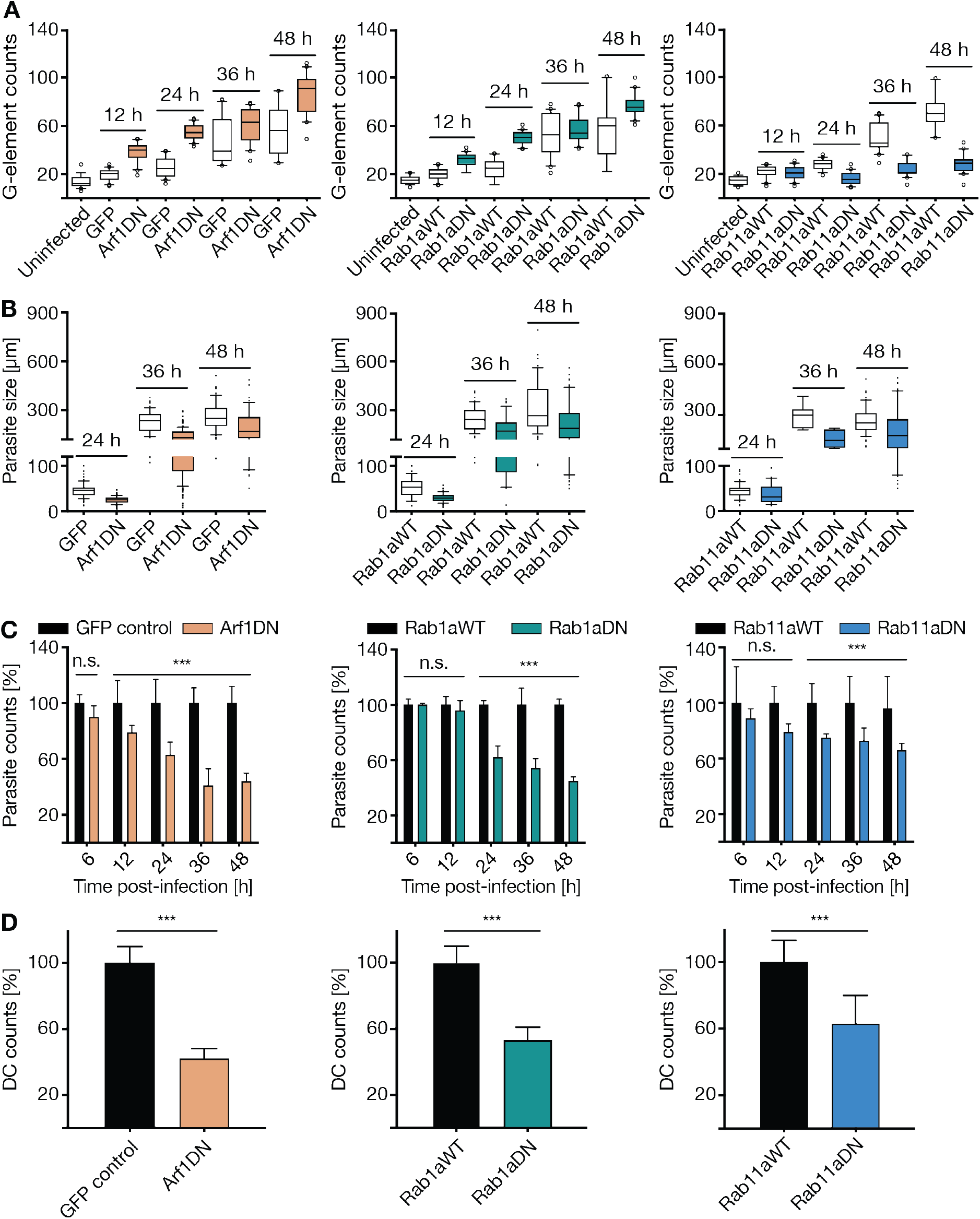
Image-based characterisation of the parasite development in host cell impaired in Golgi-associated vesicle transport. Further analysed were the three dominant negative mutants Arf1, Rab1a and Rab11a. **A)** The effect of the respective small GTPase mutant on the parasite-induced hcGolgi fragmentation was analysed based on number of G-elements/cell. Cells overexpressing either cytosolic GFP or the respective WT form of Rab1a or Rab11a served as controls. Expression of GFP or WT constructs had no effect Golgi morphology (uninfected). While Arf1DN (left panel) and Rab1aDN (middle panel) enhance the number of G-element compared to the infected control cells, Rab11aDN (right panel) blocks parasite-induced hcGolgi fragmentation. **B)** Parasite growth in Arf1DN (left panel), Rab1aDN (middle panel) and Rab11aDN (right panel) cells was assessed based on parasite sizes at 24, 36 and 48 hpi. **C)** Quantification of total parasite numbers in DN cells during the first 48 h of infection. Numbers are shown as a percentage of the control (100%) at each time point. Parasite survival is significantly affected in all three DN mutants. By 48 hpi, only >40% (Arf1DN, Rab1aDN) and >70% (Rab11aDN) parasites survive. **D)** Quantifications of detached cell (DC) at 65 hpi. In the DN cells only >40% (Arf1DN), >55% (Rab1DN) and 60% (Rab11aDN) parasites complete the liver stage by forming DCs. Percent DCs relative to GFP control. All data shown corresponds to triplicate experiments and triplicate repeats per experiment, with n = 200 per time point and transgenic cell line. *** *p* ≤ 0.01, paired *t*-test or ANOVA.

In order to dissect the impact that Arf1DN and Rab1aDN have on *Plasmodium* liver stage development, we measured parasite size (Fig. 6B, left and middle panel) and counted total number of parasites at the indicated time points (Fig. 6C, left and middle panel) and detached cells (Fig. 6D left and middle panel). The intercepted vesicle transport in the host cell affected the parasite development in two aspects. On one hand we observed a delay in parasite growth, which arises early in the 24 h old schizonts. On the other hand we detected a gradual decrease in parasite numbers (Fig. 6B-C, left and middle panel). By 48 hpi only 50 % of parasite survived (Fig. 6C, left and middle panel). The decreased survival rate is also reflected by the reduction in detached cells at 65 hpi. Relative to the infected control cells, only 40% (Arf1DN) to 55% (Rab1aDN) of parasites complete the liver stage (Fig. 6D, left and middle panel). The onset of the parasite death, however, started earlier (at 12 hpi) in the Arf1 DN cells (Fig. 6B left panel). Likewise, we observed different dynamics in hcGolgi disintegration. The number of G-elements in Arf1DN mutants rose faster than in the Rab1aDN background (Fig. 6A left panel).

Rab11a regulates the vesicular transport at the *trans* face of the Golgi and coordinates post-Golgi traffic and recycling of endosomes to the cell surface (Reviewed in (Goud et al., 2018; Welz et al., 2014)). Moreover, previous work has shown that Rab11 disruption has only little effect on hcGolgi architecture (Rejman Lipinski et al., 2009; Welz et al., 2014) (Fig. S3). When we infected cells overexpressing Rab11aDN, confocal images confirmed the manifestation of the hcGolgi at the PVM. In contrast to the other mutants, the number of G-elements per cell in the Rab11aDN background remained consistent throughout infection, showing no increase of hcGolgi fragmentation (Fig. 6A, right panel). Loss of Rab11a function prevents the parasite-induced fragmentation of the hcGolgi. Thus, Rab11a activity seems critical for the parasites ability to remodel the hcGolgi.

We continued by analysing the parasite development in context of Rab11aDN. Parasite size measurements, showed a significant growth delay in late schizonts (36-48 hpi) (Fig. 6B, right panel). The similar trend was observed regarding parasite numbers. Although parasite survival rate progressively decreased, the effect was less pronounced than in the Arf1DN and Rab1aDN mutants. About 70% parasites developed into late schizonts (48 hpi) and overall 60% formed detached cells at 65 hpi (Fig. 6B-6D, right panels). Therefore, loss of Rab11a function affects predominantly parasite proliferation. At the current state, we cannot distinguish if the impaired exo- and endocytic pathway of the host cell or the Rab11aDN-mediated parasite deficiency to induced Golgi fragmentation are the main cause of the reduction of parasite fitness in the liver.

## Discussion

The liver stages of *Plasmodium* display one of the fastest growth rates among eukaryotic cells. Nevertheless, little is known on mechanisms of nutrient scavenging or host cell subversion by the parasite, which support such massive growth. Following invasion and the formation of the parasitophorous vacuole, *Plasmodium* preferentially settle at the juxtanuclear region of the hepatocyte and shortly afterwards associate with the host cell ER (Bano et al., 2007; Kaiser et al., 2016). Although *Plasmodium* interactions with the host cell ER have been investigated, the dynamics and relevance of the hcGolgi-*Plasmodium* interactions remained largely unexplored. In the present work, we characterize the parasite relationship with hcGolgi throughout the pre-erythrocytic stage.

We observed an association of the hcGolgi with the PVM within 24 h after invasion. The PVM-hcGolgi interplay was sustained during the entire pre-erythrocytic development and accompanied by fragmentation of the hcGolgi into miniaturized complexes. Critical for the parasite-induced fragmentation of the hcGolgi is an active Rab11a, a small GTPase required for vesicular transport at the TGN of the host cell. hcGolgi fragmentation is certainly not unique to *Plasmodium.* Bacteria such as *Chlamydia* (Heuer et al., 2009; Rejman Lipinski et al., 2009)*, Salmonella*, and *Legionella* (Reviewed in (Allgood and Neunuebel, 2018; Brumell and Scidmore, 2007)), as well as various viruses (Reviewed in ((Ravindran et al., 2016; Spriggs et al., 2019)), and the parasite *T. gondii* (Romano et al., 2017) all induce hcGolgi fragmentation by mechanisms including molecular mimicry and modification of hcGolgi proteins, amongst which Golgins and Rab GTPases are common targets. In *P. berghei,* fragmentation becomes pronounced only from 24 h onwards corresponding with the initiation of parasite schizogony. At this point, the hcGolgi fragments distribute around the PVM, while in many cases losing the perinuclear localization. Importantly, the parasite-induced fragmentation of the hcGolgi preserves hcGolgi cellular function.

Previous work on *T. gondii* has shown that a preferential localization of the parasite near the microtubule organizing centre (MTOC)-Golgi region, which is the intersection of endocytic and exocytic pathways, may facilitate the interception of vesicular trafficking ensuring acquisition of lipids such as cholesterol deriving from lysosomes, or ceramide deriving from the hcGolgi (Bisanz et al., 2006; Coppens et al., 2006; Melo and Souza, 1996). Although our approach does not give insight into metabolites scavenged by *P. berghei*, we show that the parasite-hcGolgi interaction plays a critical role for a productive passage through the liver. Extrinsic interference with host cell MT or hcGolgi-associated vesicular transport impaired pre-erythrocytic development and reduced the number of detached cells. Nutrients scavenging is a well-established role of the PVM. Parasite-encoded PVM proteins have been shown to directly interact with lipid binding proteins and to sequester host lipids by hijacking endo-lysosomal vesicles (Petersen et al., 2017; Sá E Cunha et al., 2017). An alternative route represents the transport of host cell nutrients through pore and channels in the PVM (Bano et al., 2007; Mesén-Ramirez et al., 2019).

Since *T. gondii* localizes not only to the periphery of the hcGolgi, but also near the host microtubule organizing centre (MTOC) (Coppens et al., 2006; Romano et al., 2013; Walker et al., 2008), Romano *et al.* investigated whether hijacking and fragmentation of the hcGolgi by *T. gondii* might in fact be preceded by hijacking of the MTOC. Their studies in polarized and non-polarized cells, however, showed that the *T. gondii* PV first associates with the hcGolgi, and only later with the MTOC (Romano et al., 2013). We investigated whether inducing hcGolgi ribbon disassembly and vesiculation by interfering with microtubule polymerization would cause complete or partial dissociation of the hcGolgi from the PVM. The effect of each drug on microtubule conformation was consistent with previous observations (van de Moortele, 1993). Both drugs induced hcGolgi disassembly and dispersion across the cytoplasm in a reversible manner. Upon washout of the drugs the Golgi re-distributed as a fragmented structure around the parasite’s PVM, rather than re-stacking as a single Golgi ribbon in the nuclear periphery of the host cell. Interestingly, even after prolonged incubation with drugs individual G-elements did not fully detach from the PVM. The drug-resistant G-elements seem to form stable, MT-independent attachment with PVM. Similar intimate contacts were previously described for host cell ER with the PVM (Kaiser et al., 2016), suggesting the presence of membrane contact sites. The nature and cellular function of the Golgi-PVM attachment will be subject of future investigations.

Crucial for establishing the hcGolgi-parasite interaction is the presence of a functional PVM. Mutants missing the PVM-resident proteins UIS3 or UIS4 are surrounded by a functionally compromised PVM (Mueller et al., 2005a, 2005b). Both PVM-deficient parasites are not able to recruit or maintain a stable connection with hcGolgi. Engaged in a more temporal relation with the PVM and its dynamic TVN extension are host cell vesicles of the endo-lysosomal component of the autophagy pathway. This carefully controlled association guarantees optimal nutrient acquisition and prevents simultaneously the elimination of parasite (Agop-Nersesian et al., 2018, 2017; Grützke et al., 2014; Niklaus et al., 2019; Petersen et al., 2017; Prado et al., 2015; Real et al., 2018; Thieleke-Matos et al., 2016; Wacker et al., 2017). With the hcGolgi, however, parasites build a persistent relationship. Loss of the hcGolgi contact is accompanied with morphological features of parasite death suggesting that the interaction plays also a role beyond nutrient scavenging.

Strategies to overcome host defences developed by pathogens, include molecular mimicry (Galán and Collmer, 1999; Galán and Wolf-Watz, 2006; Spanò and Galán, 2018); inhibition of signal transduction cascades involved in pattern recognition receptor (PRR) activation (Espinosa and Alfano, 2004; Mogensen, 2009); inhibition of antigen presentation; or inhibition of vesicular transport through the host’s secretory pathway, with the purpose of abrogating chemokine or cytokine secretion (Burnaevskiy et al., 2013; Dong et al., 2012; Selyunin and Alto, 2011). Cargo in the secretory pathway usually follows a route including the ER, the ER Golgi intermediate compartment (ERGIC) and the Golgi. ARF and Rab family GTPases are essential in cargo packaging and delivery between compartments, and not surprisingly, various pathogens particularly target members of the RAB and ARF families to exploit nutrient hijacking. We chose to investigate the role of Arf1 on *Plasmodium* development, as well as representative Rab GTPases with functions at various points of the trafficking pathway (depicted in Fig. 1 and summarized in Table 1).

Arf1 is a member of the ADP-ribosylation factor (Arf) family, which belongs to the Ras superfamily of small GTPases. Arf1 alone, is a multi-functional component. Together with Arf2 and Arf3, Arf1 is thought to be key in regulating assembly of different types of coat complexes on budding vesicles along the secretory pathway and to activate lipid-modifying enzymes (Bonifacino and Glick, 2004; Singh et al., 2019). Among its functions, Arf1 plays a key role in the early secretory pathway, primarily with regards to retrograde transport from the Golgi to the ER and between Golgi cisternae via recruitment of the coat protein complex I (COPI) to budding transport vesicles (Bonifacino and Glick, 2004; Singh et al., 2019). Amongst various candidates tested, overexpression of Arf1DN was the most detrimental for *Plasmodium* development, with deleterious effects on parasite sizes and numbers detected since very early on in infection. Despite equally successful sporozoite invasion rates as in WT cells, parasites show poor survival, development, and ultimately relatively unsuccessful completion of the pre-erythrocytic cycle. Indeed, the downregulation of vesicular coat subunits (COPB2 and COPG1) in the COPI complex have also shown to impair *Plasmodium* development in the liver (Raphemot et al., 2019). Moreover, the β-COP subunit vesicles to also localize to the PVM in late schizonts. Various pathogens have previously been shown to target Arf1, including *Legionella pneumophila* (Reviewed in (Isberg et al., 2009)) and *E. coli* (Selyunin et al., 2014). *Legionella* activates and recruits Arf1 to establish vesicular trafficking of the host’s Golgi to the phagosome, providing the latter with ER-like characteristics which favour bacterial development (Amor et al, 2005; Nagai et al., 2002). Like-wise, *E. coli* simultaneously inhibits Arf1 and Rab1, while Arf1-binding-deficient mutants are unable to disrupt Golgi architecture (Selyunin et al., 2014).

Regarding the Rab proteins tested, candidates included Rab1a and Rab2, both of which localize to the ER/*cis*-Golgi region and are involved in ER-Golgi transport; Rab6a, which localizes to the Golgi and is involved in retrograde Golgi traffic; and Rab11a, which localizes to the recycling endosomes and the *trans-*Golgi network. Among these, the most striking effects on *Plasmodium* development were observed in Rab1a, and Rab11aDN cells, while overexpression of dominant negative forms of Rab2a and Rab6a had nor overall impact on parasite fitness. Although *Plasmodium* itself induces Golgi fragmentation, and we hypothesize this is beneficial for nutrient acquisition and for increasing surface area contact sites with the PVM, artificial disassembly of the host cell Golgi either by microtubule manipulation or by Arf1 or Rab1 dominant negative overexpression is detrimental for the parasite. We hypothesize that the type of fragmentation induced by the parasite ensures a functional Golgi, while allowing maximal nutrient hijacking. Conversely, Rab1a or Arf1DN overexpression dismantle the Golgi complex and result in a non-functional Golgi. Regarding Rab11 observations, previous work on *Chlamydia trachomatis* had shown that both Rab6 and 11 are important regulators of infection, that their disruption prevented *Chlamydia*-induced Golgi fragmentation, and that this resulted in inhibition of transport of nutrients to *Chlamydia* inclusions (Rejman Lipinski et al., 2009). In our work, we observed similarly, reduced hcGolgi fragmentation upon dominant negative overexpression of Rab11a. In contrast to Rab6aDN expression, Rab11aDN showed significant effects on parasite development, suggesting that functions specific to Rab11, beyond hcGolgi fragmentation are important. Rab11a localizes to recycling endosomes, and is key for anterograde trafficking, from the trans-Golgi network to the plasma membrane. Its inhibition has previously been shown to result in reduced surface delivery of cargo while cargo trafficking from ER to Golgi and intra-Golgi remains unaffected. Together with the clathrin adaptor protein GGA1, Rab11a represents an additional host cell factor involved in the *trans*-Golgi to endosome vesicular transport (Raphemot et al., 2019).

With host-targeted therapies becoming a new line of intervention, understanding the nature of the host-parasite interaction is of great importance. In this study, we not only describe the dynamics and structural changes of the hcGolgi upon infection but also shed some light on the importance of the hcGolgi in *Plasmodium* pre-erythrocytic development. We envisage that future work will give further interesting insights into the role, the hcGolgi might play in nutrient scavenging and parasite survival.

## Materials and Methods

### Ethics statement

Animal studies were carried out under the approval of the Animal Research Ethics Committee of the Canton Bern, Switzerland (Permit Number: 91/11 and 81/11); and the University of Bern Animal Care and Use Committee, Switzerland. For all studies, Balb/c females 5-8 weeks of age, weighing 20 to 30 g at the time of infection were used. Mice were bred in the central animal facility of the University of Bern or supplied by Harlan Laboratories. Blood feeding of mosquitoes was performed under Ketasol/Dorbene anaesthesia, and all efforts were made to minimize suffering.

### *Plasmodium berghei* lines

Various *P. berghei*-ANKA lines were used to infect mice and cultured cells. Transgenic lines used in this study included Pb^Hsp70^mCherry (referred to as PbmCherry), constitutively expressing mCherry in the parasite cytosol (Burda et al., 2015); Pb ^Hsp70^mCherry ^ef1α^NLuc (referred to as PbNLuc), constitutively expressing mCherry and the luminescent protein NanoLuc (De Niz et al., 2016); Pb^Lisp2^Exp1-mCherry (Graewe et al., 2011) which has mCherry-tagged Exp1 under the liver stage-specific Lisp2 promoter, enabling visualization of the PVM from 36 hpi onwards; and Pb^UIS4^ UIS4-mCherry (Grützke et al., 2014), which expresses the mCherry-tagged PVM protein UIS4 under its endogenous promoter and enables PVM visualization from early times post-infection. We used also the parasite mutants Pb ΔUIS4 and PbΔUIS3 (Mueller et al., 2005b, 2005a).

### Parasite maintenance in mosquitoes

Balb/c mice were treated with phenylhydrazine 3 days prior to intra-peritoneal infection with the different *P. berghei* lines. After three days of infection, parasites were spotted on a coverslip, and exflagellation was determined. The infected mice were then used to feed 100-150 *Anopheles stephensi* female mosquitoes each. Mice were anaesthetized with a combination of Ketasol (125 mg/kg) and Dorbene (12.5 mg/kg) anaesthesia, and euthanized with CO_2_ after completion of the feed. Afterwards, mosquitoes were fed until use with 8% fructose and 0.2% Para-aminobenzoic acid (PABA). Sporozoites used for hepatocyte infection were isolated form the mosquitoes’ salivary glands between 16 and 26 days post-feed.

### Mammalian cell culture and infections

The human hepatoma HepG2 cell lines (European Collection of Cell Culture) and the human epithelial HeLa cell lines (kind gift from Robert Menard) were maintained in minimum essential medium (MEM) containing Earle’s salts supplemented with 10% heat-inactivated fetal calf serum (FCS), 2mM L-Glutamine, and 100 U Penicillin, 100 μg/ml Streptomycin (cMEM) (all from PAA laboratories, E15-024, A15-101, P11-010, M11-004). Cells were kept at 37°C in a 5% CO_2_ cell incubator and were split every 4 d by treatment with accutase. For infections, 20,000 cells were seeded into 24-well plates and infected with *P. berghei* sporozoites with an MOI of 1. Infected cells were maintained in cMEM and 2.5 μg/ml amphotericin B (AT-MEM) (PAA laboratories, P11-001) to avoid fungal contamination. AT-MEM was exchanged every 12 h. Selected time points for assaying infection were 6, 12, 24, 36, 48, and 56 hpi.

### Plasmids

The plasmid pEGFPN1-GalT (plasmid 11929) provided by Jennifer Lippincott-Schwartz (Cole et al., 1996) and the plasmid pcDNA HA Arf1 DN T31 N (plasmid 10833) provided by Thomas Roberts (Furman, et al; 2002) were obtained from Addgene. Plasmids pECFP-C1-Rab1a WT, pECFP-C1-Rab1a DN (AS mutation), pECFP-C1-Rab2 WT, pECFP-C1-Rab2 DN, pECFP-C1-Rab6a WT, pECFP-C1 Rab6a DN, pECFP-C1-Rab11a WT and pECFP-C1-Rab11a DN provided by Won Do Heo (Department of Biological Sciences KAIST, Korea). Plasmid EGFP-CLASP1 DN overexpressing the microtubule-binding domain was a kind gift from Helder Maiato (Universidade do Porto, Portugal).

### Transient transfection of mammalian cells

For each transfection 5× 10^5^ cells were re-suspended in 100 μl filter-sterilized transfection buffer containing 120 mM Na-Phosphatase, 5 mM KCl, 20 mM MgCl_2_, 5 mM NaCl, pH 7.2 and mixed with 2.5 μg of the respective plasmid. Cells were electroporated using the T-028 program of a Nucleofactor™ 2b transfection device (Lonza). For the IFA analysis approximately 1/10^th^ of the transfected cells were seeded in each well of a 24 well-plate, and glass bottom dishes (MatTek) for live cell imaging were prepared with 1/6^th^ of the total transfection.

### Treatment with microtubule-targeting drugs Nocodazole and Taxol

Following infections with Pb^Hsp70^mCherry sporozoites, cells were treated with 50 nM Nocodazole (M1404, Sigma-Aldrich), or 50 nM Taxol (T7402, Sigma-Aldrich) for 1 h at 24 hpi. For recovery experiments, cells were washed 3 times with AT-MEM, and fixed after 15 min or 1 h, respectively. For analysis of parasite development, infected cells were treated for 1 h with 50 nM Nocodazole or Taxol, respectively at three different time points 12 hpi, 24 hpi and 36 hpi. After treatment, cells were washed 3 times with AT-MEM, and incubated again on AT-MEM until fixation with 4% PFA. Cells were fixed at 48 hpi, after which parasite numbers and sizes in wells were imaged and quantified using the image analysis software Fiji (URL: http://fiji.sc/Fiji). Additionally, at 65 hpi, detached cells were quantified. Control cells for each time point were treated with the drug solvent, DMSO.

### Immunofluorescence assays

Infected HepG2 or HeLa cells were fixed at the indicated time points in 4% paraformaldehyde (PFA) in phosphate-buffered saline (PBS, 137 mM NaCl (Sigma-Aldrich, S9888-1Kg), 2.7 mM KCl (Fluka Chemie AG P9541-1kg), 10 mM Na_2_HPO_4_ (Sigma-Aldrich, S5136-500g), 1.8 mM KH2PO4 (Sigma-Aldrich, P5655), pH 7.4) for 10 min and subsequently permeabilized with 0.1% Triton X-100 in PBS (Fluka Chemie, T8787-250ml), for another 10 min. After 1 h blocking with 10% FCS-PBS cells were stained with the diluted primary antibodies for an additional hour. For fluorescent labelling, cells were subsequently incubated with secondary antibodies for 1 h and nuclei visualized with 1 μg/ml DAPI (Invitrogen, D-1306). All incubations were performed at room temperature. For microscopy, cells were mounted on microscope slides using Dako Fluorescent Mounting Medium (Dako, S3023).

The primary antibodies used were mouse monoclonal anti-GFP (Roche, 11814460001), rabbit polyclonal anti-GFP (Acris, SP3005P), rabbit polyclonal anti-UIS4 (kindly provided by P. Sinnis), chicken polyclonal anti-Exp1 (Heussler lab, Bern, Switzerland), mouse monoclonal anti-GM130 (BD-Biosciences, 610822), mouse monoclonal anti-α tubulin (Sigma, T9026), and rat monoclonal anti-HA (Roche, 11867423001). All primary antibodies were used at a dilution of 1:500 in 10% FCS/PBS. As secondary antibodies, species-specific Alexa Fluor® 488 (Invitrogen Molecular Probes, A-11001; A-11008), Alexa Fluor® 594 (Invitrogen Molecular Probes, A-21209; A-11032) or Cy5 (Dianova, 111-175-144; 703-175-155) were used at a final concentration of 0.2 μg/ml.

### Live cell imaging and time lapse microscopy

Confocal microscopy was performed on fixed and live cells with a SP8-STED Leica microscope. 3D images were acquired using a 63x oil objective (NA 1.4) and the LAS X software (Leica) used for image acquisition. Optical z-sections with 0.5 μm spacing were acquired using the LAS X software. Time lapse microscopy on live cells was performed using a Leica DMI6000B microscope equipped with a 63x water objective (NA 1.4), and the Leica Application Suite (LAS) AF software.

### Serial block face scanning electron microscopy

5×10^4^ HeLa cells were seeded in every well of a 96 well plate and infected with Pb^Hsp70^ mCherry parasites, as described above. Infected cells were FACS-sorted at 6 hpi for Pb^Hsp70^mCherry-infected cells and re-seeded at a density of 10,000 sorted cells per well into a 96 well optical plate (Greiner bio one). Following this, parasites were fixed in a glutaraldehyde buffer at 48 hpi and processed as per previously published protocols (Deerinck et al., 2010; Kaiser et al., 2016). SBF-SEM images were acquired with a Quanta FEG 250 (FEI Company) equipped with a Gatan 3View2XP ultramicrotome (accelerating voltage = 3.5 kV; low vacuum). Images were processed using Fiji.

### Bioluminescence imaging

HeLa cells transiently expressing the indicated plasmids, were infected with Pb ^Hsp70^mCherry ^ef1α^NLuc (De Niz et al., 2016) sporozoites. At different times post-infection, cells were either detached from the wells using accutase, or detached cells were collected at 65 hpi from the medium supernatant, and lysed in 20 μl of 1x passive lysis buffer (PLB) (Promega) for downstream analysis. Lysates were transferred to 96-well black plates (Greiner BioOne), and imaged using an IVIS Lumina II system. For consistent and unsaturated measurements, the optimal NanoGlo™ (Promega) assay substrate dilution used for probing liver stage parasites was 1:500. Uninfected cells were used as background controls.

### Image analysis

The shortest distance between the PVM and the host cell Golgi was measured using Imaris software on a 3D reconstructed cell volume. Distances are displayed in a box-plot and show the 10-90 percentile of the distances measured in 100 cells at each time point. All results appearing as individual dots fall over or under this remaining 10-percentile. No outliers were excluded from calculations or *p-*values.

The number and volume of GalT stacks present in *Plasmodium-*infected and uninfected cells were measured on 3D reconstructed images using Imaris software (Bitplane), and Fiji. Individual foci were identified by intensity, using automatic thresholding. Parasite sizes were quantified using Fiji, and automatic fluorescence thresholding.

### Statistical analyses

Data are displayed in box plots or histograms generated using PRISM 6.0 software. Means, medians, interquartile ranges, and SDs were calculated from 3 independent experiments performed in triplicate, using STATA 13.0 software. The *p-*values were calculated using Student’s *t* test or ANOVA in STATA 13.0.

## Acknowledgements

We are grateful to Freddy Frischknecht for critically reading the manuscript and helpful discussions. We thank Helder Maiato, Photini Sinnis, Ann-Kristin Müller, and Kai Matuschewski for sharing plasmids, reagents and parasite lines. The Microscopy Imaging Center (MIC), University of Bern provided access to the microscopes and Theodor Kocher Institute facilities at University of Bern provided access to the IVIS Lumina II system.

## Authors Contributions

M.D.N. performed all experiments unless specified otherwise. G.K. and B.Z. performed and analysed the electron microscopy experiment. W.D.H. provided the Rab-GTPase constructs. V.T.H. secured funding and provided critical input and guidance. M.D.N. and C.A.N. designed experiments, analysed, visualized and interpreted the data, and wrote the manuscript. C.A.N. conceptualized and supervised the study. All authors provided critical feedback to the final manuscript.

---

**Figure S1.**
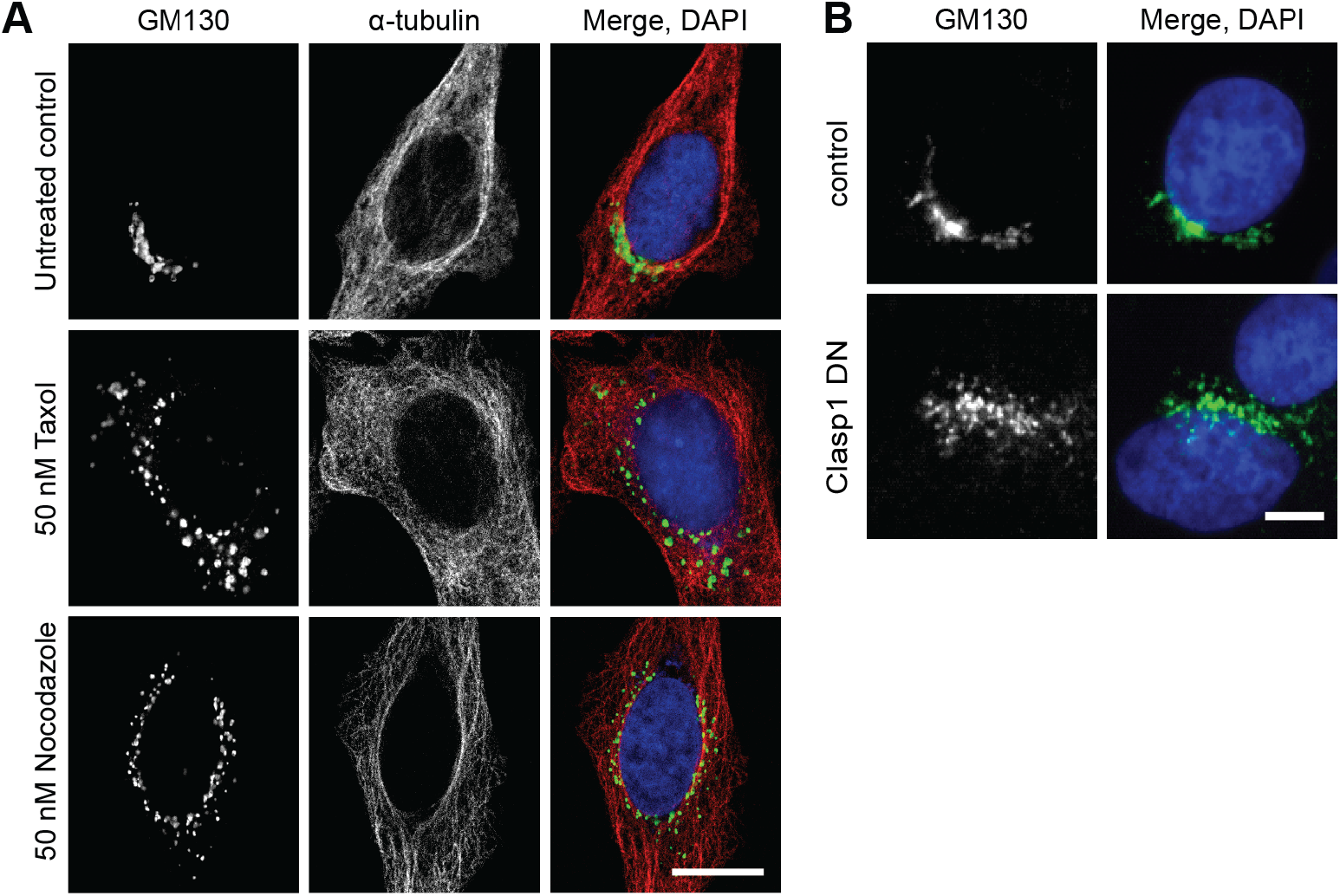
Manipulation of host cell microtubules result in a dismantled hcGolgi. **A)** HeLa cells treated with either 50 nM of Taxol or Nocodazole show hcGolgi disassembly and dispersion across host cell cytoplasm. Confocal images were taken from cells stained against the *cis*-Golgi protein GM130 (green), α-tubulin (red) and DAPI (blue). **B)** Hela cell transiently overexpressing the microtubule binding domain of the CLIP-associating protein 1 (CLASP1) in HeLa cells result in the loss of the Golgi architecture. hcGolgi of transfected cell was visualized with GM130 (green). Scale bar 10 μm.

**Figure S2.**
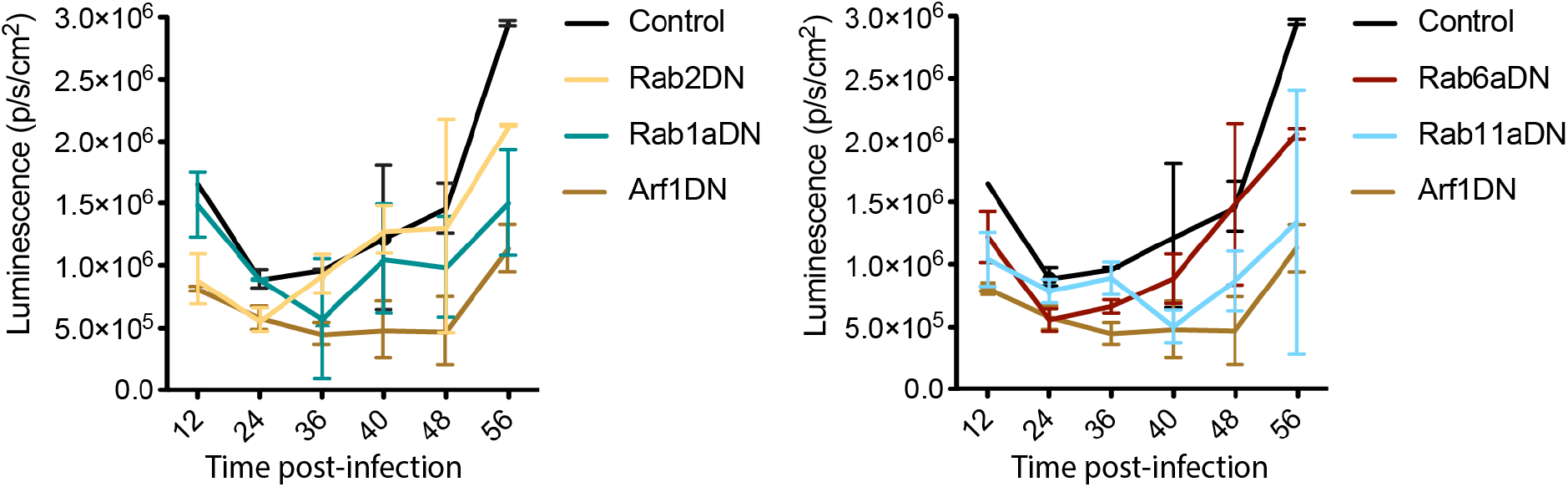
Bioluminescence analysis of the *P. berghei* liver-stage development in cell impaired for Golgi-associated vesicular transport. HeLa cells transiently overexpressing either GFP (control), or one of the dominant negative GTPase (Arf1DN, Rab1aDN, Rab2DN, Rab6aDN and Rab11aDN) were infected with NanoLuc-expressing *P. berghei* parasites (PbNLuc). Quantitative representation of luminescence measurements. Consistently, higher luminescence was detected in GFP control cells, followed by Rab6aDN, Rab2DN, Rab1aDN, Rab11aDN. The Arf1DN displayed the strongest effect on parasite development.

**Figure S3.**
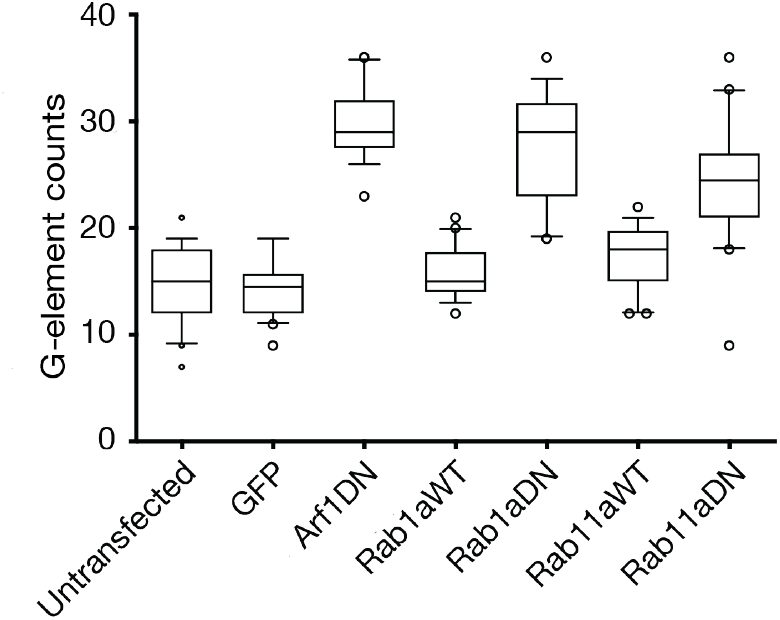
Overexpression of Golgi-associated small GTPases induce hcGolgi fragmentation. Fragmentation of the hcGolgi was assessed in HeLa cells transiently overexpressing a dominant negative version of the small GTPases Arf1, Rab1a or Rab11a. As controls served untransfected cells, cell overexpressing GFP (as reference for Arf1DN) or the wild type (WT) version of Rab1a or Rab11a, respectively.

